# Angiotensin antagonist inhibits preferential negative memory encoding via decreasing hippocampus activation and its coupling with amygdala

**DOI:** 10.1101/2021.08.14.456361

**Authors:** Ting Xu, Xinqi Zhou, Guojuan Jiao, Yixu Zeng, Weihua Zhao, Jialin Li, Fangwen Yu, Feng Zhou, Shuxia Yao, Benjamin Becker

**Author notes:** **Correspondence to** Benjamin Becker, University of Electronic Science and Technology, Xiyuan Avenue 2006, 611731 Chengdu, China, mail.

## Abstract

Exaggerated arousal and dysregulated emotion-memory interactions are key pathological dysregulations that accompany the development of post-traumatic stress disorder (PTSD). Current treatments for PTSD are of moderate efficacy and preventing the dysregulations already during exposure to threatening events may attenuate the development of PTSD-symptomatology. In a preregistered double-blind, between-subject, placebo-controlled pharmaco-fMRI design, the present proof-of-concept study examined the potential of a single dose of angiotensin II type 1 receptor (AT1R) antagonist losartan (LT) to attenuate the mnemonic advantage of threatening stimuli and the underlying neural mechanism via combining an emotional subsequent memory paradigm with LT (n=29) or placebo treatment (n=30) and a surprise memory test after 24h washout. LT generally improved memory performance and abolished emotional memory enhancement for negative yet not positive material while emotional experience during encoding remained intact. LT further suppressed hippocampus activity during encoding of subsequently remembered negative stimuli. On the network level LT reduced coupling between hippocampus and basolateral amygdala during successful memory formation of negative stimuli. Our findings suggest that LT may have the potential to attenuate memory formation for negative yet not positive information by decreasing hippocampus activity and its functional coupling strength with amygdala. These findings suggest a promising potential of LT to prevent preferential encoding and remembering of negative events, a mechanism that could prevent the emotion-memory dysregulations underlying the development of PTSD-symptomatology.

## Introduction

Memory enhancement for threatening events and stimuli facilitates the avoidance of future dangers and thus represents an important evolutionary survival mechanism (1). However dysregulation in this fine-grained interplay between emotion and memory may crucially contribute to the development of major psychiatric disorders (2), particularly post-traumatic stress disorder (PTSD) (3, 4). PTSD is a prevalent and highly debilitating psychiatric disorder (5), which is characterized by hyperarousal and re-experiencing traumatic events frequently through disruptive memories and dissociative flashbacks (6). Neurocognitive models of PTSD propose that symptoms manifesting as a consequence of trauma-induced dysregulations in emotion cognition interactions lead to biased encoding and subsequent memory for threat-associated information (3, 7, 8). While maladaptive emotion-memory integration has been considered as a trans-diagnostic treatment target (9), the current interventions aim at suppressing exaggerated emotional responses or modifying the traumatic memory following the development of a psychiatric condition (10, 11). With respect to PTSD the available treatments are characterized by modest efficacy and high drop-out rates (12), and administering pharmacological agents immediately after trauma exposure did not result in the expected reduction of PTSD symptomatology (13, 14). As such a preventive pharmacological strategy which modulates encoding during the exposure to emotional events may represent an innovative and promising approach to prevent the development of PTSD symptoms.

Growing evidence from animal models and human findings have suggested a crucial role of the renin-angiotensin system (RAS) in emotion and memory formation which may overlap with processes relevant for the development of PTSD (15–17). Different lines of research have indicated an important role of the RAS in the development of PTSD such that (1) blood levels of renin were elevated in individuals with trauma-experience (18), (2) treatment with angiotensin blockers, particularly the angiotensin II type 1 receptor antagonist losartan (LT) during trauma exposure reduced the incidence of PTSD (19, 20), and (3) administration of LT in healthy controls also reduced autonomic stress reactivity during experimental trauma induction (21) and shifted emotional reactivity from negative to positive information (22, 23). While these findings support that targeting the RAS via LT may prevent the development of PTSD by modulating emotional or memory formation processes, the underlying behavioral and neurobiological mechanisms have not been examined.

Extensive research has implicated the amygdala and hippocampus in emotional memory formation, in the development of PTSD, and as potential target sites for the modulatory effects of LT on emotion and memory. The memory advantage of emotional over neutral material strongly depends on the interaction between the amygdala and hippocampus (1), with the amygdala mediating emotional arousal effects on the hippocampal formation which primarily encodes information into long term memory (24, 25). Recent neuroimaging studies have demonstrated dysregulated hippocampal and amygdala processing during fear conditioning (26) and encoding of negative emotional stimuli (3), as well as regionally specific decreased volumes of the amygdala-hippocampal (27, 28) in PTSD patients. Treatment evaluation studies moreover have reported that functional dysregulations in amygdala and hippocampus normalize with decreasing PTSD symptomatology during successful interventions (29, 30), while these regions also mediated the modulatory effects of pharmacological agents on emotional memory formation (31). The hippocampus and amygdala exhibit dense expressions of central angiotensin receptors that are considered to critically mediate learning-related neuroplasticity in these regions (32). Animal models further support this by demonstrating that LT-induced enhanced general memory function and reduced contextual memories are mediated by effects on the hippocampus (15, 33), while effects on fear-related processes are primarily mediated by the amygdala (16, 34). Initial pharmacological neuroimaging studies in humans also demonstrated LT-induced modulation of the amygdala and its connections with prefrontal regulatory regions during threat processing (17, 35). Together these animal and human studies may indicate a potential neural pathway which may mediate the attenuation of LT administration to negative memory formation.

Against this background the present preregistered randomized double-blind placebo-controlled pharmaco-fMRI experiment sought to provide proof-of-concept for the treatment potential of LT in PTSD by examining the behavioral and neural effects of a single-dose of 50mg LT on emotional memory encoding and subsequent recognition. Following treatment administration participants were presented with negative, positive and neutral pictures during fMRI and underwent a surprise memory test after a washout period of 24 hours (similar approach see (36)). Based on previous findings suggesting an LT-induced facilitation of aversive memory extinction (15, 37, 38), we expected that LT administration would particularly abolish the memory enhancement effects for negative stimuli. We further hypothesized these effects would be mediated by decreased amygdala or hippocampal activity during encoding reflecting LT-induced attenuated emotional arousal or attenuated memory formation for negative materials, respectively. Given the critical role of interactions between the hippocampus and the amygdala for emotional memory formation, specifically the memory advantage of negative events, we expected that treatment-induced changes in regional activity would be mirrored in the functional interaction between these regions.

## Methods and Materials

### Participants

N=66 healthy male participants were recruited for the present pre-registered randomized double-blind between-group pharmacological fMRI experiment. To account for variance related to sex differences in emotional memory related neural activity (39, 40) and the response to RA blockade (41) only male participants were enrolled in the present study. A total of 7 participants were excluded and lead to a final sample size of n=59 for further analysis (LT, n=29, placebo (PLC), n=30). For general enrollment criteria as well as exclusion criteria see Supplementary Methods: Participants and CONSORT flow diagram.

The study procedures adhered to the latest reversion of the Declaration of Helsinki and were approved by the local ethics committee. All participants provided written informed consent for this study which was preregistered on Clinical Trials.gov (https://clinicaltrials.gov/ct2/show/NCT04606225).

### Study protocols

A pharmacological fMRI experiment with a double-blind randomized, placebo-controlled, between-subject design was utilized to examine LT effects on encoding emotional information and subsequent memory performance. Prior to fMRI experiment participants were randomly assigned to administer either a single 50-mg p.o. dose of LT or PLC packed in identical capsules. An independent researcher not in direct contact with participants and experimenters dispensed the capsules according to a computer-generated randomization list. Psychometric assessments regarding mood, attention and memory capacity were administered at baseline and post experiment, whereas cardiovascular activity (i.e., blood pressure, heart rate) were measured at baseline, after drug administration and the experiment (Details see Supplementary Methods). LT crosses the blood brain barrier and its peak plasma levels are reached after 90 minutes while the terminal elimination half-live of LT ranges between 1.5-2.5h (42–45). The experiments thus started during peak plasma and the memory test was scheduled after 24h washout. Participants first performed a reward processing task (Duration: 30 minutes) and next underwent the emotional memory paradigm during fMRI. Results of the reward processing task will be reported in a separate publication. After the experiments, participants were asked to guess whether they had received LT or PLC. 24 hours after the experiment a computerized surprise memory recognition test was implemented (**Figure 1A**).

**Fig. 1.**
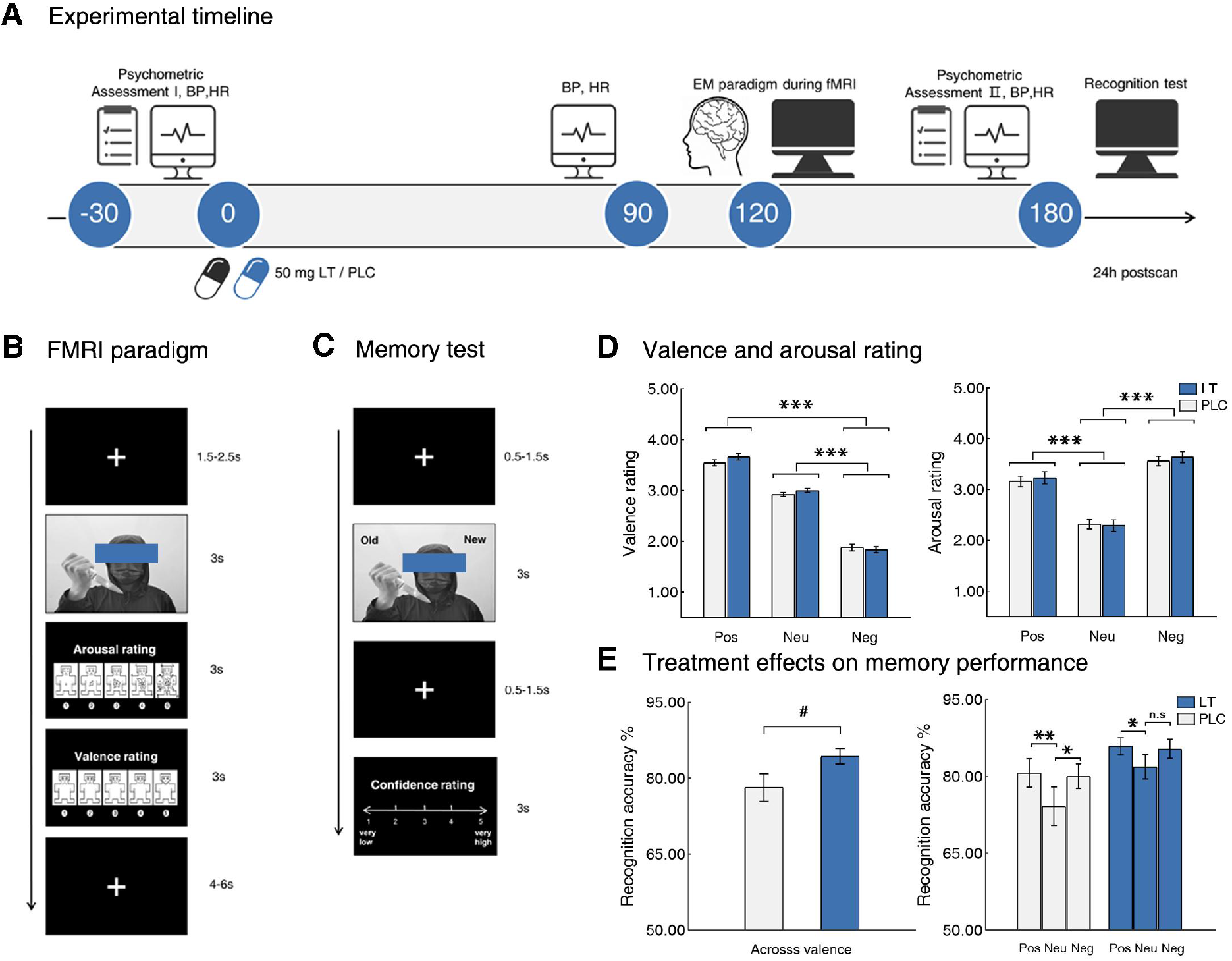
Experimental timeline, experimental paradigm and treatment effects on emotional encoding and subsequent memory performance. **A** Experimental protocol. **B-C** Schematic representation and timing of the emotional memory and recognition paradigm. **D** Emotional characteristics of the paradigm, including a valence-dependent effect on valence and a negative-specific effect on arousal. **E** Treatment effects on memory performance, indicating that LT enhanced general memory whereas it abolished the memory-enhancing effects of negative emotional stimuli. BP-blood pressure; HR-heart rate; LT-losartan; PLC-placebo; EM-emotional memory; Pos-positive; Neg-negative; Neu-neutral; n.s-not significant, #p<0.08, *p<0.05, **p<0.01, ***p<0.001.

### Experimental paradigm

The event-related fMRI emotional memory paradigm incorporated positive, neutral and negative pictorial stimuli from the Nencki Affective Picture System (46) and International Affective Picture System (47). Three runs with 10 pictures per emotional condition were used and the order was balanced between treatment conditions (Stimuli details see **Supplementary Methods**). Each stimulus was presented for 3s during encoding, followed by a jittered interval ranging from 1.5s to 2.5s and separate ratings for valence and arousal by means of the Self-Assessment Manikin scales (ranging from 1 (minimum) to 5 (maximum)) (48). The rating scale was presented for 3s and presentation order was balanced across participants. The trials were separated by a jittered inter-trial interval of 4-6s which served as low-level baseline.

24 hours after encoding participants completed a forced choice recognition memory test in the laboratory. The test included the 90 pictures previously shown during encoding intermixed with 54 novel images (18 per emotional condition, matched for valence and arousal; see **Supplementary Methods**). During the surprise memory test participants had to indicate whether the stimuli had been shown during encoding (‘old’ or ‘new’) and to rate their confidence for the response (**Figure 1B-C, Supplementary Methods**).

### Behavioral data analyses

Statistical analyses were conducted in JASP (version 0.14.1.0). Effects of LT on emotional experience during encoding (valence, arousal) were analyzed by means of mixed ANOVAs with treatment (LT, PLC) as the between-subjects factor and valence (positive, negative, neutral) as the within-subjects factor. In line with previous studies (49, 50) recognition accuracy was computed by subtracting the false alarm rate for each emotional valence from its respective hit rate. Emotion-specific effects of LT on subsequent memory performance were next investigated by means of a mixed ANOVA with treatment (LT, PLC) as the between-subjects factor, valence (positive, negative, neutral) as the within-subjects factor and recognition accuracy as the dependent variable. To facilitate a more sensitive and specific evaluation of LT’s effect on the emotional enhancement of memory we further examined the difference in emotion memory effect (DEm) by comparing recognition accuracy for each emotional condition with the neutral condition within each treatment group (in line with previous works (51, 52)). To control for the potential influence of arousal differences between the valence categories on the valence-specific treatment effects, we recomputed the corresponding analyses including arousal as covariate and results remained stable (see **Supplementary Methods**).

### fMRI data acquisition, preprocessing and analysis

MRI data were collected on a 3.0 Tesla system (GE MR750, General Electric Medical System, Milwaukee, WI, USA) and further preprocessed using standard procedures in SPM 12 (Statistical Parametric Mapping; http://www.fil.ion.ucl.ac.uk/spm/;Wellcome Trust Centre for Neuroimaging) (Details provided in **Supplementary Methods**). We modelled two separate event-related general linear models (GLM) on the first level to separately examine effects of LT on emotional experience during encoding and subsequent memory formation.

The first GLM model examined effects of treatment on emotional experience and modelled presentation of each stimulus according to the valence condition (positive, negative, neutral) using an event-related design and convolution with the hemodynamic response function (53). Rating periods (arousal, valence) and six head motion parameters were included as covariates. In line with our a priori regional hypotheses the group level analysis focused on the amygdala and hippocampus as defined by Jülich-Düsseldorf brain (JuBrain) cytoarchitectonic atlas based structural masks (54, 55). The extracted estimates from hippocampus and amygdala were subjected to two separate mixed ANOVA models with treatment as the between-subject factor and valence as the within-subject factor. In line with the behavioral analysis a second GLM was designed to facilitate a sensitive examination of the emotional memory effect (DEm). To this end the DEm was modelled by contrasting the remembered positive or negative items respectively with the non-remembered items (similar approach see (56)). On the group level the main DEm effect was examined across emotions and treatment groups (“positive + negative remembered items>all forgotten items”) within the structurally defined amygdala-hippocampal complex to identify regions engaged in emotional memory formation irrespective of a treatment or emotional condition. Effects of LT modulation on the positive and negative DEm effects were subsequently examined by mixed ANOVA models with treatment as between-subjects variable and valence as within-subjects variable on the extracted parameter estimates from independent atlases-based masks of the identified regions. All analyses incorporated adequate correction for multiple comparisons, on the voxel-level family-wise error (FWE) correction adopted to the size of the region-of-interest and a threshold value of p<0.05 was employed (details see **Supplementary Methods**).

To additionally explore network level effects of treatment a generalized form of context-dependent psychophysiological interaction analysis was conducted. Based on the BOLD level results the right hippocampus (as defined by an independent atlas mask) was chosen as a seed region. We modeled condition-dependent variations in functional connectivity from right hippocampus to each voxel in the whole brain. In line with our a priori hypothesis on hippocampus-amygdala functional coupling the analyses focused on the anatomically defined amygdala (threshold at PSVC-FWEpeak<0.05 within the bilateral amygdala mask) (details see **Supplementary Methods**). To control the potential confounding effects of arousal, we repeated all fMRI analyses based on the extracted parameters estimates after including arousal of all valence as covariates and observed consistent results (see **Supplementary Methods**).

## Results

### Sociodemographic, psychometric and physiological characteristics

The LT (n=29) and PLC (n=30) groups exhibited comparable sociodemographic and psychometric properties (**Table 1**;all ps>0.13). Previous studies suggested that the effects of LT on cardiovascular activity was only observed after at least 3 hours while the central effects occurred in a faster manner (57, 58). Consistent with this, 80 minutes after the LT application examination of blood pressure and heart rate revealed no significant treatment difference (all ps>0.11, **Supplementary Results**,**Table S1**). Mood and anxiety assessments before the 24h memory assessments did not reveal significant group differences (all ps>0.24, see **Supplementary Results**,**Table S2**).

**Table 1.** Sociodemographic and Psychometric assessments

|  | Time | PLC (n=29) | LT (n=30) | t test, p value |
| --- | --- | --- | --- | --- |
| Age, years |  | 20.83±2.05 | 20.48±2.98 | 0.52 |
| Body Mass Index, kg/m2 |  | 21.19±2.17 | 21.28±2.01 | 0.86 |
| PANAS - positive | Baseline | 26.80±5.89 | 27.38±4.86 | 0.68 |
|  | Post experiment | 25.87±6.19 | 24.69±5.44 | 0.44 |
| PANAS - negative | Baseline | 15.07±5.12 | 17.03±7.48 | 0.24 |
|  | Post experiment | 13.17±5.08 | 14.45±6.60 | 0.41 |
| STAI - state | Baseline | 39.77±8.86 | 42.28±10.32 | 0.32 |
|  | Post experiment | 38.17±7.78 | 40.41±9.46 | 0.32 |
| STAI - trait | Baseline | 41.13±7.19 | 41.90±9.85 | 0.73 |
|  | Post experiment | 40.27±7.81 | 41.59±8.83 | 0.55 |
| BDI II | Baseline | 8.43±8.97 | 9.17±7.07 | 0.73 |
|  | Post experiment | 6.17±7.31 | 7.34±6.01 | 0.50 |
| DCAT1 | Baseline | 38.13±5.71 | 38.17±6.56 | 0.98 |
|  | Post experiment | 56.17±9.06 | 55.55±8.02 | 0.78 |
| DCAT2 | Baseline | 65.10±12.91 | 64.17±10.59 | 0.76 |
|  | Post experiment | 47.07±6.24 | 45.28±6.52 | 0.29 |
| DCAT3 | Baseline | 61.50±9.76 | 58.72±8.33 | 0.25 |
|  | Post experiment | 70.50±12.56 | 68.55±9.15 | 0.50 |
| Working memory | Baseline | 0.86±0.09 | 0.86±0.09 | 0.87 |
|  | Post experiment | 0.91±0.08 | 0.87±0.10 | 0.14 |

Values are presented as mean ± SD. PLC, placebo; LT, Losartan; PANAS, Positive and Negative Affect Schedule; STAI, Spielberger State-Trait Anxiety Inventory; BDI II, Beck Depression Inventory II; DCAT 1-3, Digit cancellation test 1-3.

### Behavioral results

The mixed ANOVA with treatment (LT, PLC) as the between-subjects factor, valence (positive, negative, neutral) as the within-subjects factor, and valence or arousal ratings as dependent variable revealed no significant main or interaction effect involving treatment (all ps>0.29), but a significant main effects of valence (valence, Greenhouse-Geisser, F_(1.43,57)_=542.02, p<0.001, partial η^2^=0.91; arousal, F_(2,57)_=227.06, p<0.001, partial η^2^=0.80, **Figure 1D**). The neutral pictures were rated less positive than positive ones (t_(58)_=12.01, p<0.001, d=1.56), and were rated less negative than negative ones (t_(58)_=20.54, p<0.001, d=2.68). The negative pictures were perceived as more arousing than positive (t_(58)_=6.50, p<0.001, d=0.85) and neutral pictures (t_(58)_=20.83, p<0.001, d=2.71) irrespective of treatment.

Examining LT effects on subsequent recognition accuracy by means of a treatment (LT, PLC) × valence (positive, negative, neutral) mixed ANOVA revealed no significant interaction effect (p>0.66), but a significant main effect of valence (Greenhouse-Geisser, F_(1.77,57)_=6.12, p<0.01, partial η^2^=0.10) and a marginally significant main effect of treatment (F_(1,57)_=3.77, p=0.06, partial η^2^=0.06). Post-hoc tests revealed an enhanced recognition performance for emotional relative to neural pictures in both groups (positive vs. neutral, t_(58)_=3.18, p<0.01, d=0.41; negative vs. neutral, t_(58)_=2.85, p=0.01, d=0.37), and a general improved memory performance across valence following LT administration (t_(57)_=1.94, p=0.06, d=0.25, **Figure 1E**).

To test our hypothesis that LT will specifically attenuate memory formation for negative events we directly examined DEm within each treatment group. In the PLC group recognition memory for negative (t_(29)_=2.10, p<0.05, d=0.38) and positive pictures (t_(29)_=2.81, p<0.01, d=0.51) was significantly enhanced relative to neutral ones, whereas the LT group exhibited a memory-enhancing effects only for positive stimuli (t_(28)_=2.34, p<0.05, d=0.43, **Figure 1E**) yet not for negative stimuli (t_(28)_=1.35, p=0.19, d=0.25, **Figure 1E**). However, the direct group comparison failed to reach significance (ps>0.05).

### LT decreases hippocampus activity during encoding for negative stimuli

Examination of extracted parameter estimates from the amygdala revealed no significant main or interaction effect of treatment (all ps>0.12), but a significant main effect of valence (F_(2,57)_=9.53, p<0.001, partial η^2^=0.14, **Figure 2A**). The post-hoc tests indicated higher amygdala activity during encoding of negative pictures relative to neutral (t_(58)_=4.28, p<0.001, d=0.56) and positive ones (t_(58)_=2.90, p<0.01, d=0.38) in both treatment groups. In contrast a significant valence × treatment interaction effect (F_(2,57)_=3.69, p=0.03, partial η^2^=0.06) was observed in the hippocampus, whereas no main effects of valence or treatment (all ps>0.14) were observed in this region. Post-hoc tests revealed that following LT hippocampus activity was significantly decreased during encoding for negative stimuli compared to both positive (t_(28)_=-2.96, p<0.01, d=-0.55, **Figure 2B**) and neutral ones (t_(28)_=-2.43, p<0.05, d=-0.45, **Figure 2B**).

**Fig. 2.**
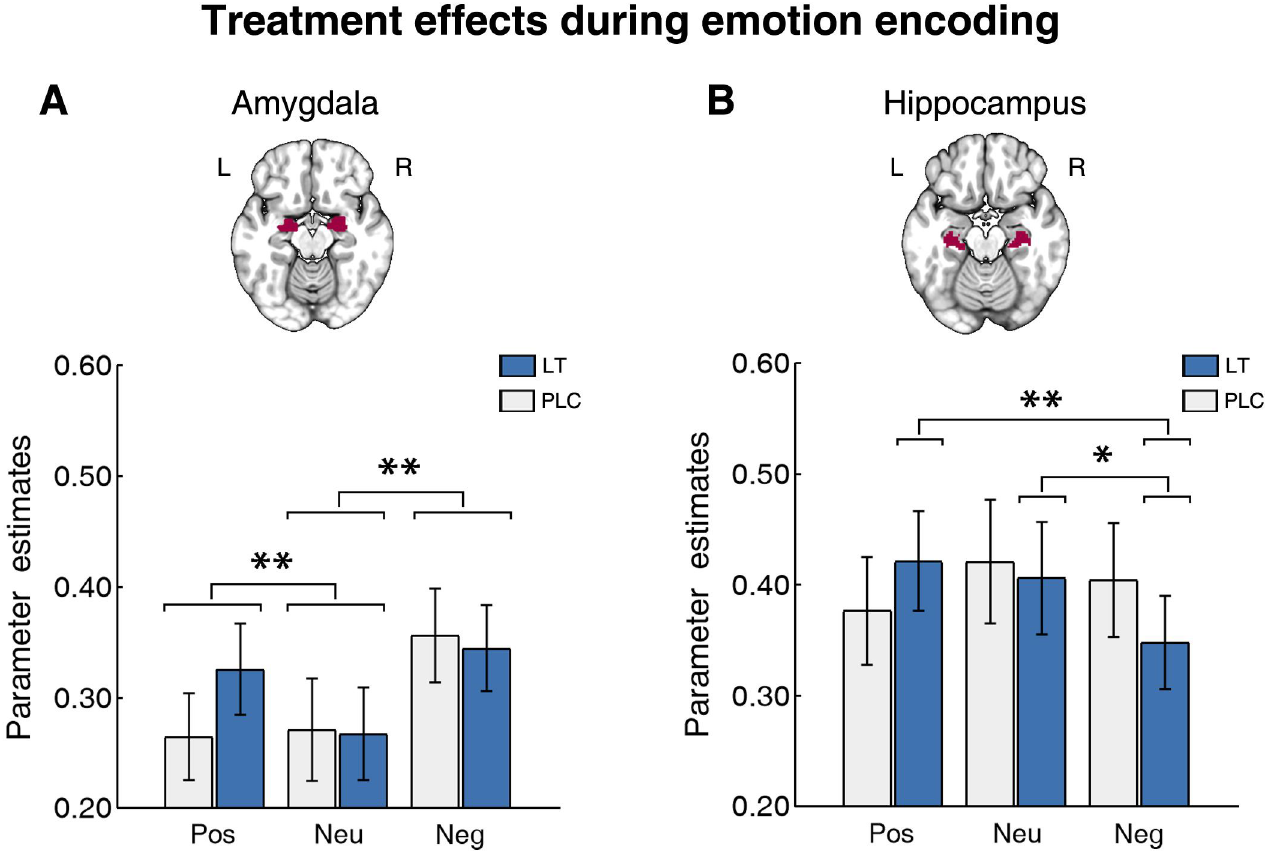
Examination of LT effects on neural activation during emotion encoding. **A** The amygdala showed increased activity during encoding for negative stimuli relative to positive and neutral ones across treatment. **B** Following LT hippocampus activity decreased significantly when encoding negative stimuli compared to the positive and neutral ones whereas there was no difference for hippocampus response in PLC group. Error bars represent standard error of the mean. Pos-positive; Neg-negative; Neu-neutral; PLC-placebo; LT-losartan; *p<0.05,**p<0.01.

### LT decreases hippocampus activity during subsequent memory formation for negative stimuli

In general the successful emotional memory formation was associated with increased responses in the bilateral amygdala (left, peak MNI, −20/-2/-28, T_(53)_=4.84, P_SVC-FWEpeak_=0.002, k=180; right, peak MNI, 22/-2/-28, T_(53)_=4.05, P_SVC-FWEpeak_=0.02, k=119, **Figure S2**) and the right hippocampus (peak MNI, 30/-28/-16, T_(53)_=4.24, P_SVC-FWEpeak_=0.01, k=21, **Figure S2**).

In a next step effects of treatment on the DEm in the identified amygdala-hippocampal nodes involved in successful memory formation were examined by means of extraction of parameter estimates from independently defined atlas-based regions for the amygdala and hippocampus. Examination of the amygdala revealed a significant main effect of valence (left, F_(1,52)_=19.49, p<0.001, partial η^2^=0.27; right, F_(1,52)_=10.62, p<0.01, partial η^2^=0.17), but no main or interaction effect involving treatment (all ps>0.22). Post-hoc tests demonstrated increased amygdala activity for subsequent memory formation for negative pictures relative to positive ones (left, t_(54)_=4.42, p<0.001, d=0.60; right, t_(54)_=3.26, p<0.01, d=0.44,**Figure S3**). The right hippocampus only exhibited a marginal valence × treatment effect (F_(1,52)_=3.33, p=0.07, partial η^2^=0.06). Specifically, this region displayed comparable activity during subsequent memory formation for positive and negative stimuli during PLC (t_(26)_=-0.30, p=0.77, d=-0.06), while during LT decreased activity for negative memory formation compared with positive memory formation (t_(26)_=-2.49, p=0.02, d=-0.48, **Figure 3A**).

**Fig. 3.**
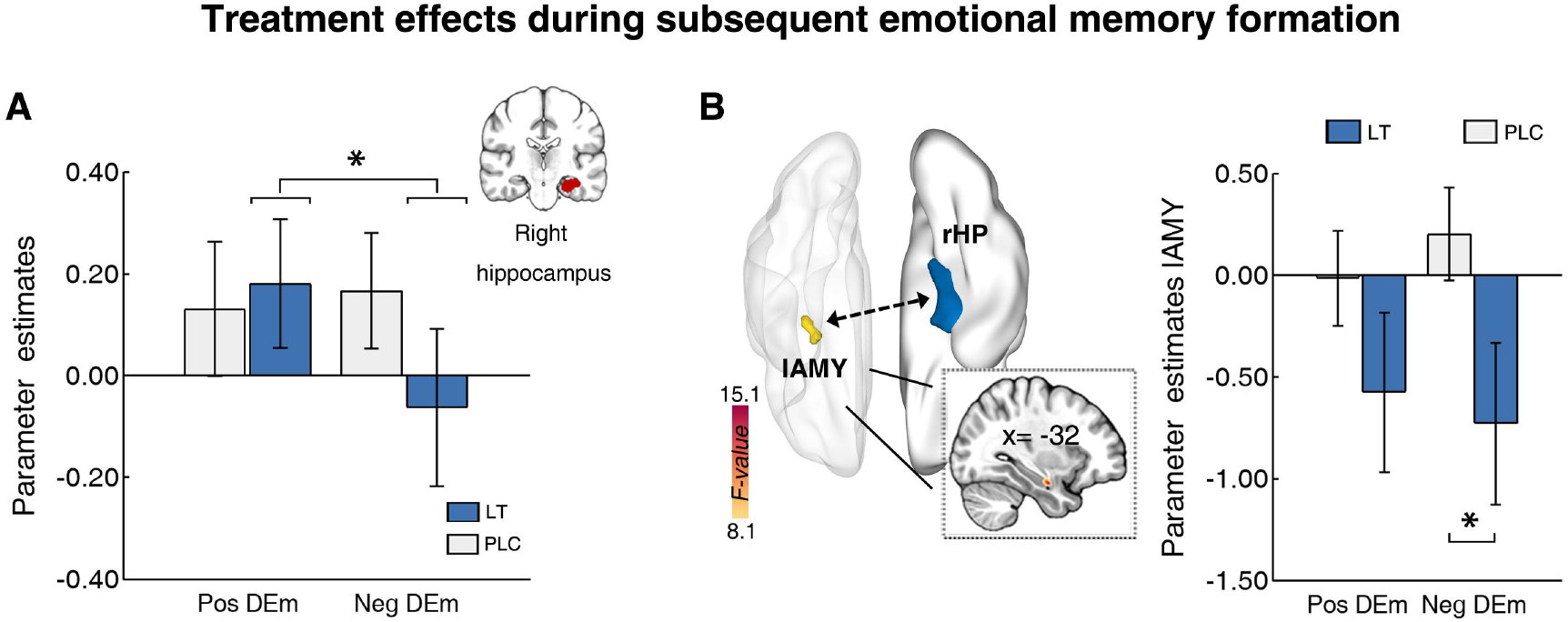
LT effects on neural activity and connectivity during successful memory formation for emotional stimuli (DEm effect). **A** Right hippocampus activity decreased during memory formation for negative stimuli following LT administration while comparable hippocampus activation was observed for successful encoding of positive and negative stimuli following PLC. **B** Interaction effects of valence with treatment was observed in the functional connectivity between seed region right hippocampus and left amygdala. LT specifically decreased coupling strength of the right hippocampus with left basolateral amygdala during memory formation of subsequently successfully remembered negative stimuli. Error bars represent standard error of the mean. lAMY, left amygdala; rHP, right hippocampus; Pos, positive; Neg, negative; PLC, placebo; LT, losartan. *p<0.05.

### LT decreases hippocampal-amygdala coupling during subsequent memory formation for negative stimuli

Given the importance of hippocampus-amygdala interaction for emotional memory formation a functional connectivity analysis was conducted to examine treatment effects on the coupling between these regions. During successful emotional memory formation a significant valence × treatment interaction effect was found on right hippocampus coupling with the left amygdala (peak MNI, −32/-6/-20, P_SVC-FWEpeak_=0.02, k=8, F_(1,52)_=14.87, **Figure 3B**). Probabilistic mapping demonstrated that the identified cluster was located in the basolateral amygdala (BLA) subregion (78.1% probability). Extracting parameter estimates from the left amygdala revealed that LT significantly decreased right hippocampus coupling strength with left amygdala during successful memory for negative stimuli relative to PLC (t_(52)_=-2.04, p<0.05, d=-0.55).

## Discussion

The present study examined whether pharmacological modulation of the RAS via LT during emotional encoding can attenuate the subsequent memory advantage of negative material and whether this effect is mediated by effects on the amygdala or hippocampus. In line with our prediction, LT slightly enhanced general memory performance with tentative evidence suggesting that this was accompanied by an inhibition of the memory advantage for negative information in the absence of modulatory effects on emotional arousal or valence. On the neural level the effects were accompanied by decreased hippocampus activity specifically during encoding of negative stimuli, while amygdala activation was not affected. On the network level LT specifically decreased the functional communication of the right hippocampus with the left BLA during successful memory for negative stimuli. Together these findings suggest a contribution of the RAS to memory formation, particularly for negative stimuli, via valence-specific modulation of hippocampus activation and its coupling with the BLA. In contrast, emotional evaluation and associated amygdala activity during encoding remained unaffected by RAS blockade.

Despite converging evidence emphasizing the role of hyperarousal and a negative memory encoding bias during trauma exposure for the development of PTSD, the underlying mechanisms have been difficult to target therapeutically (3, 7, 8). Previous observational studies demonstrated that blockade of the RAS via LT during trauma exposure reduced the incidence of PTSD and hypothesized that the beneficial effects may be mediated by an LT-induced attenuation of autonomic reactivity or arousal (19, 20). In the present study LT did not affect subjective arousal experience while it slightly enhanced memory performance. Although following LT the general memory performance showed a trend to significant improvement, which is in accord with previous findings (59, 60), LT additionally induced a valence-dependent shift in emotional memory formation. In particular, when focusing on the DEm effect (reflecting the relative memory advantage of emotional relative to neutral material) the LT-treated group maintained a memory advantage of positive (relative to neutral) pictures but abolished the relative memory advantage of negative pictures. Our findings tentatively suggest that LT may have the potential to attenuate preferential memory formation for negative stimuli via an indirect arousal-mediated mechanism but rather a valence-specific reduction in the memory advantage of negative emotional material. Although the direct group comparison did not reach statistical significance the behavioral functional profile of LT could point to a potential that reduce exaggerated emotional memory formation for negative materials, a mechanism which may attenuate the risk for memory dysregulations that facilitate the development of PTSD.

Neuroimaging studies in PTSD commonly linked symptoms in the domains of hyperarousal and emotional memory to dysregulations in the amygdala-hippocampal complex. Some evidence suggests that hyperarousal and emotional-reactivity symptoms in PTSD are strongly mediated by amygdalar dysregulations (61, 62) while memory dysfunctions are associated with hippocampal dysregulations (63). However, during threat learning and encoding of negative emotional materials both regions have been found to be dysregulated in PTSD (26, 52, 64, 65). The present study demonstrated that LT specifically decreased hippocampus activity during encoding of negative stimuli while not affecting amygdala functioning. This finding may explain the behavioral effects of an LT-induced abolishment of memory advantage of negative material in the absence of effects on arousal. Moreover animal models have demonstrated an important role of the hippocampus in mediating effects of LT on aversive memory, such that (1) an LT-induced reduction in fear memories triggered by interoceptive cues was mediated by attenuated cFOS expression in the hippocampus (33), and (2) the preference for safe relative to threat environments was mediated by an LT-induced suppression to hippocampal AT1R (66).

In addition to the hippocampus, the amygdala plays a critical role in emotion-memory integration (1) as well as emotional memory dysregulations in PTSD (52). The amygdala is considered as a key locus for enhanced responses to emotionally arousing (25), particularly threatening stimuli (67) and consequently modulates the arousal-mediated enhancement of emotional memories. Although a previous study reported an LT-induced prolonged amygdala threat reactivity in humans (35), the present study did not observe direct effects on amygdala activity yet a valence-specific attenuated functional coupling between the right hippocampus and the BLA during negative memory formation. Whereas LT may not have affected arousal or emotional processing associated with amygdala activity per se, it suppressed communication between the amygdala and the hippocampus - a pathway that has been strongly involved in encoding, consolidation and reconsolidation for negative and threatening materials across species(68, 69).

Previous studies also explored pharmacological strategies to attenuate emotional memory formation, e.g. by means of the β-adrenergic blocker propranolol, the NMDA receptor antagonist ketamine, or 3,4-Methylenedioxymethamphetamine. However, the effects were commonly mediated by biased emotion-associated amygdala reactivity (31), impaired general memory performance (51), as well as abolished emotional enhancement for both positive and negative materials (70). In contrast, LT (1) specifically abolished the advantage of negative material, while not impairing general or positive memory performance (e.g. NMDA antagonists), and (2) did not affect general valence or arousal processing which may lead to biased reactions or decision making. Together, the pattern of observed behavioral and neural effects of LT may point to a promising mechanism to prevent exaggerated negative emotional memories already during the exposure stage while preserving general memory formation and emotional processing.

Finally, our results have to be interpreted in the context of limitations. First while the present study reduced variance related to gender difference future works should confirm the generalization of the observed effects to female individuals. Second, given the relatively short retention interval of 24h longer term effects of LT on emotional memory and consolidation need to be determined. Third, although the wash-out period between encoding and memory assessment allowed us to control for acute treatment effects on recognition future studies may additionally monitor interim factors that may affect memory performance such as sleep quality or stress exposure (71, 72). Further, emotional stimuli can initiate complex interactions between hormonal systems that are coordinated by the hypothalamic–pituitary –adrenal axis (HPA) encompassing central and peripheral sites (73). Previous animal models also suggest a contribution of both central and peripheral AT1R to HPA mediated emotional processes (74). As such the potential contribution of effects on peripheral AT1R to the central effects of LT needs to be explored in future studies. Finally, future studies may also implement paradigms that allow the application of complex network level analyses such as ‘background connectivity’ which has been associated with consolidation of memory after 24 hour (75).

In conclusion, the present study provides the first evidence for a role of the RAS in valence-specific memory formation via modulating the hippocampus and its connections with the BLA. RAS blockade via LT abolished the memory advantage for negative events in the absence of effects on emotional processing. On the neural level this attenuation was mirrored in valence-specific decreased hippocampus activity and coupling of this region with the BLA. Together these findings provide proof-of-concept for the functional profile of LT to inhibit dysregulated negative memory formation, a mechanism which have the potential to prevent the development of trauma-induced emotional memory dysregulations.

## Acknowledgements and disclosure

This work was supported by the National Key Research and Development Program of China (2018YFA0701400). Data availiability: unthresholded group-level statistical maps are available on OSF (https://osf.io/v4j9d/). Other data of this study is available from the corresponding author upon reasonable request. The authors report no biomedical financial interests or potential conflicts of interest.

## Author contributions

TX and BB designed the study; TX, GJ, YZ, XZ conducted the experiment and collected the data. TX, XZ, FZ, FY, LJ performed the data analysis. TX and BB wrote the manuscript draft, WZ and YS critically revised the manuscript draft.

