## supplementary for "Angiotensin antagonist inhibits preferential negative memory encoding via decreasing hippocampus activation and its coupling with amygdala"

### *Supplementary information*

### Supplementary Methods

#### Participants: sample size calculation and exclusion criteria

From the N = 66 participants initially enrolled in the study data from n = 1 was excluded due to technical failure of the MRI system, n=6 were excluded as outliers in terms of the primary behavioral outcome (i.e. recognition performance beyond three standard deviations from the respective group mean) leading to a total of n=59 participants for the final analysis (Losartan = 29, Placebo = 30). See also **Figure S1** showing the CONSORT flow diagram. A priori sample size calculation based on G-power (Faul et al., 2007) indicated that this sample size was sufficient to determine interaction effects ($\eta^2 = 0.25$) that could detect a treatment effects with acceptable power of 0.95 (at $\alpha = 0.05$). All participants were free from current or a history of psychiatric, neurological, or other medical disorders. Additional exclusion criteria encompassed: excessive head movement (>3 mm translation or 3° rotation), current or regular use of psychotropic substances including nicotine, a body mass index <18 or >24.9, visual or motor impairments, and contraindications for MRI or losartan (LT).

#### Assessment of potential confounders

To control for pre-treatment between group differences in emotional state and unspecific effects of treatment on emotional state the Positive and Negative Affective Schedule (PANAS), Spielberger State-Trait Anxiety Inventory (STAI) and the Beck Depression Inventory (BDI-II) were administered before drug administration and after the experiment. To control for potential effects of treatment on cardiovascular activity heart rate and blood pressure were assessed before and after drug administration, as well as after the MRI acquisition. Working memory capacity was also assessed via completing a 2-back letter working memory task before drug administration and after the MRI experiment. To control for potential confounding effects of LT on attention, all participants finished a digit cancellation test (D-CAT) before treatment administration and after the experiment.

### Experimental paradigm

We adopted the emotional memory paradigm from a previous pharmaco-fMRI study (Becker et al., 2017). A total of 90 positive, negative and neutral pictures from the International Affective Picture System (Lang et al., 1997) and Nencki Affective Picture System (Marchewka et al., 2014) were used in the fMRI paradigm. Pictures were presented in three separate runs with 10 pictures of each valence (30 pictures per run). Subjects were instructed to give arousal and valence ratings towards each picture stimuli. The arousal and valence scores of pictures in separate runs were balanced: mean valence (SD): Run1, positive 6.98 (0.46), negative 2.73 (0.42), neutral 5.28 (0.42), Run2, positive 7.10 (0.48), negative 2.64 (0.36), neutral 5.25 (0.48), Run3, positive 7.08 (0.72), negative 2.77 (0.29), neutral 5.38 (0.45); mean arousal (SD), Run1, positive 6.36 (0.41), negative 6.30 (0.35), neutral 4.21 (0.37), Run2, positive 6.38 (0.36), negative 6.25 (0.35), neutral 4.25 (0.32), Run3, positive 6.17 (0.48), negative 6.15 (0.44), neutral 4.16 (0.36), all ps>0.48, one-way ANOVA comparing valence or arousal ratings between runs. Mean arousal for positive and negative pictures in each run also showed no difference, all ps>0.50, paired samples t-test.

Similar to the procedure reported by Striepens (Striepens et al., 2012), at 24 hours postscan, subsequent memory performance was tested by a force choice recognition memory test. During recognition task, participants were shown the 90 target pictures previously presented in the scanner along with 54 novel pictures. These novel stimuli were also matched for emotional arousal and valence between runs; Run1, positive 6.95 (0.20), negative 2.78 (0.53), neutral 5.31 (0.23), Run2, positive 6.99 (0.58), negative 2.81 (0.29), neutral 5.27 (0.50), Run3, positive 7.06 (0.54), negative 2.94 (0.22), neutral 5.40 (0.37); mean arousal (SD), Run1, positive 6.09 (0.58), negative 6.47 (1.09), neutral 4.22(0.19), Run2, positive 6.18 (0.54), negative 6.23 (0.18), neutral 4.32 (0.28), Run3, positive 6.22 (0.55), negative 6.33 (0.50), neutral 4.51 (0.30), all ps>0.61, one-way ANOVA comparing valence or arousal ratings between runs. Positive and negative distracted pictures in each run also showed no difference for the emotional arousal rating, all ps>0.51, paired samples t-test. Participants were not told how many pictures they saw during picture presentation, therefore no expectation of the amount of to be recognized pictures was mentioned to the subjects. Each picture was presented for 3s followed by a fixation screen for 500ms to 1500ms and participants had to decide via button press whether each picture had been shown during the scanning (‘old’) or not (‘new’). After picture presentation participants had 3s to rate their confidence for the decision on a five-point Likert scale (1-very low confident, 2-low confident, 3-unsure, 4-confident, 5-highly confident). The mixed ANOVA with treatment (LT, Placebo) as the between-subject factor, valence (positive, negative, neutral) as the within-subject factor, and confidence as dependent variable revealed no interaction effect (Greenhouse-Geisser, $F_{(1.76, 57)}$=1.91, p=0.15, partial $\eta^2$=0.03), but a marginal significant main effects of treatment ($F_{(1.76, 57)}$=3.66, p=0.06, partial $\eta^2$=0.06) and a significant main effects of valence ($F_{(1.76, 57)}$=17.75, p<0.001, partial $\eta^2$=0.24). The marginal main effect of treatment reflects a trend for a higher confidence in the LT group and aligns with a similar treatment effect pattern observed for memory accuracy. Post-hoc tests on the valence main effect revealed that negative pictures were recalled with more confidence than positive ($t_{(58)}$=4.14, p<0.001, d=0.54) and neutral pictures ($t_{(58)}$=5.01, p<0.001, d=0.65) irrespective of treatment.

#### MRI data acquisition, preprocessing and analysis

MRI data were collected on a 3.0 Tesla system (GE MR750, General Electric Medical System, Milwaukee, WI, USA) using a standard 12 channel heal coil. The high-resolution brain anatomical MRI image were acquired using a T1-weighted sequence (TR = 6 ms; TE = 2 ms; flip angle = 9°; field of view = 256 × 256 mm; matrix size = 256 × 256; voxel size = 1 × 1 × 1 mm; number of slices, 156; slice thickness, 1 mm) to improve spatial normalization of the functional data and exclude subjects with apparent brain pathologies. Functional data using blood oxygenation level-dependent (BOLD) contrast were obtained on this scanner using a T2*-weighted echo planar imaging sequence (TR = 2000 ms; TE = 30 ms; slices number, 39; slice thickness, 3 mm; flip angle = 90°; field of view = 240 × 240 mm; voxel size = 3.75 × 3.75 × 4 mm; resolution = 64 × 64).

All fMRI images were preprocessed and analyzed using the standard procedure in SPM 12 (Statistical Parametric Mapping; http://www.fil.ion.ucl.ac.uk/spm/;Wellcome Trust Centre for Neuroimaging). The first 5 volumes of each functional time series were discarded to allow for T1 equilibration. Remaining images were corrected for acquisition time delay, realigned to correct for head motion, unwarped for magnetic field inhomogeneities correction, and co-registered with the T1-weighted structural image. To further control for the impact of motion we computed mean framewise displacement (FD) which indexes the mean volume-to-volume head motion (Power et al., 2012; Power et al., 2014). The mean FD was significantly less than 0.5mm (Mean ± SEM, 0.35 ± 0.02,

$t_{(58)}$ = -9.81, $p < 0.001$) across all runs and was comparable between the LT and placebo groups (LT, 0.32 ± 0.02; Placebo, 0.37 ± 0.03, $t_{(57)}$ = 1.60, p = 0.12). In addition we also extracted the temporal component based noise correction variant (tCompCor) by conducting CompCor analysis in fMRIPrep (Esteban et al., 2019) to estimate physiological noise components (Behzadi et al., 2007) and then included these into a group comparison analysis. The averaged component estimates across all runs were comparable between the treatment groups (all ps>0.05, see **Table S3**), arguing against confounding effects of strong effects of treatment induced physiological effects. A fast diffeomorphic registration algorithm (Diffeomorphic Anatomical Registration using Exponentiated Lie Algebra, DARTEL) was further used for segmentation to account for inverse consistent deformation in image registration. After that the images were normalized to Montreal Neurological Institute (MNI) standard space (interpolated to 2 × 2 × 2 mm voxel size) using the segmentation parameters from the anatomical images, and were then spatially smoothed using an isotropic Gaussian kernel with full-width at half-maximum (FWHM) of 8 mm.

We modelled two separate event-related general linear models (GLM) on the first level to separately examine effects of LT on emotional experience during encoding and subsequent memory formation.

The first GLM model for examining effects of treatment on emotional experience included presentation periods of each emotional picture separated by valence (positive, negative, neutral) that were modeled for each run using a stick function convolved with the standard hemodynamic response function (Friston et al., 1994). The rating periods (arousal, valence) and six head motion parameters were included as covariates. In line with our a priori regional hypotheses the group level analysis focused on the amygdala and hippocampus as defined by Jülich-Düsseldorf brain (JuBrain) cytoarchitectonic atlas based structural masks (Amunts et al., 2005; Amunts & Zilles, 2015). The extracted estimates from hippocampus and amygdala were subjected to a mixed ANOVA with treatment as the between-subject factor and valence as the within-subject factor.

The second GLM specifically aimed at determining effects of treatment on emotional memory formation and therefore modelled neural activity reflecting the difference in memory (Dm) effect by differentiating subsequently remembered from subsequently forgotten items (Paller & Wagner, 2002). For the Dm analysis items that were subsequently remembered and forgotten were modelled into separate regressors with valence/arousal rating-periods and head motion parameters being included as covariates. To increase the sensitivity of the analysis and in line with previous studies on memory and emotional memory formation (Dolcos et al., 2005; Yu et al., 2012) only items that were recognized with high confidence were defined as 'remembered' (confidence = 5). Given that the total number of forgotten items was rather low and pooled over the emotional conditions, and thus the emotional-specific Dm effect was estimated as emotion-specific remembered items minus all forgotten items (similar approach see REFERENCE Becker et al., 2017, NeuroImage). Nevertheless, data from n=5 participants had to be excluded due to the absence of forgotten items (LT, n=3; PLC, n=2). In line with the behavioral analyses the contrasts for the positive and negative Dm effect were modelled on the individual level and subjected to second-level analysis. In an initial step the main effects of Dm was examined across emotions and treatment groups ("positive + negative remembered items > all forgotten items") within the structurally defined amygdala-hippocampal complex to identify regions engaged in emotional memory formation irrespective of a treatment or emotional condition bias. Effects of LT modulation on the positive and negative Dm effects were subsequently examined by mixed ANOVA models with treatment as between-subjects variable and valence as within-subjects variable on the extracted parameter estimates from the identified regions.

**Controlling effects of arousal**

In the present stimulus data set the negative stimuli induced a stronger arousal in the participants irrespective of treatment (see also **Figure 1D in the main text**). Given that previous studies reported an influence of arousal on emotional memory formation (Kensinger, 2004; Kensinger & Corkin, 2004; Tambini et al., 2017) we re-ran analyses that indicated valence-specific effects of LT including arousal as a covariate and the results remained stable. Specifically on the behavioral level, we first directly examined difference in emotion memory effect (DEm) within each treatment group after controlling effects of arousal. Here we used four linear models with recognition accuracy of emotional pictures as the dependent variable while recognition accuracy of neutral pictures was included as independent variable and the corresponding arousal was the controlled variable. We found that in placebo group, recognition memory for negative ($\beta=0.41$, $t=4.76$, $p<0.001$, 95% confidence interval [CI], [0.23,0.58]) and positive ($\beta=0.56$, $t=6.53$, $p<0.001$, 95% CI, [0.38,0.74]) pictures was significantly enhanced relative to neutral ones, while following LT administration enhanced memory was observed only for positive ($\beta=0.52$, $t=4.75$, $p<0.001$, 95% CI, [0.29,0.61]) yet not the negative pictures (β=0.25, t=1.49, p=0.15, 95% CI, [-0.09,0.59]). On the neural level, we recomputed the effects of LT on hippocampus and amygdala activity during emotion encoding using two separate mixed ANOVA models with arousal as the covariate. In line with the original analyses examination of the amygdala revealed no significant main or interaction effect of treatment (all ps>0.12), while the hippocampus exhibited a significant valence × treatment interaction effect ($F_{(2, 54)}$=3.65, p=0.03, partial $\eta^2$=0.06). Next, with respect to treatment effects on brain regions identified during subsequent memory formation (i.e., left and right amygdala, right hippocampus) we also found a robust result after controlling for arousal. Consistent with the original analyses only the right hippocampus exhibited a marginal valence × treatment effect ($F_{(1, 50)}$=3.48, p=0.07, partial $\eta^2$=0.07), while the bilateral amygdala was not affected (all ps>0.22). Likewise, the connectivity analyses examining effects of treatment on hippocampal-amygdala coupling still revealed a significant valence × treatment interaction effect ($F_{(1, 50)}$=5.77, p=0.02, partial $\eta^2$=0.10) after controlling arousal. Together these findings indicate that the valence-associated effects of LT cannot be fully explained by the different levels of arousal induced by the stimuli.

### Supplementary Results

#### Effects of LT on cardiovascular indices

Cardiovascular assessments indicated no significant treatment effects on blood pressure (diastolic or systolic) or heart rate throughout the experimental timeline (all ps>0.10, see **Table S1**).

**Table S1. Cardiovascular activity measures**

| Physiological Measures | Baseline | | Peak plasma (90min) | | Post experiment | | F test, p value |
|---|---|---|---|---|---|---|---|
| | Placebo | Losartan | Placebo | Losartan | Placebo | Losartan | |
| Systolic blood pressure | 117.73±6.46 | 120.24±7.35 | 115.73±6.62 | 115.52±9.48 | 119.20±5.57 | 119.55±6.82 | 0.12 |
| Diastolic blood pressure | 70.30±6.71 | 70.07±7.12 | 68.83±7.01 | 69.31±7.18 | 71.77±6.37 | 69.59±6.38 | 0.16 |
| Heart rate | 75.27±10.36 | 75.20±11.94 | 69.63±7.98 | 67.34±8.60 | 68.93±9.84 | 68.10±9.04 | 0.57 |

Values are presented as mean±SD.
*F* tests for the interaction of treatment (Placebo, Losartan) × time (Baseline, Peak plasma (90min), Post experiment).

**Table S2**. **Mood and anxiety assessments before the 24h memory assessments**

| | Placebo(n=29) | Losartan(n=30) | t test, p value |
|---|---|---|---|
| PANAS - positive | 26.23±6.13 | 25.17±4.83 | 0.47 |
| PANAS - negative | 13.17±5.46 | 14.21±6.14 | 0.49 |
| STAI - state | 38.20±8.1 | 40.90±9.71 | 0.25 |
| STAI - trait | 40.13±7.60 | 41.31±8.76 | 0.58 |
| BDI Ⅱ | 6.43±7.51 | 7.90±6.81 | 0.44 |

Values are presented as mean ± SD.

PANAS, Positive and Negative Affect Schedule; STAI, Spielberger State-Trait Anxiety Inventory; BDI II, Beck Depression Inventory II.

**Table S3. Group comparison on physiological noise components**

| Component | Groups | Mean±SD | t |
|---|---|---|---|
| tCompCor1 | Placebo | 0.14±0.12 | 0.70 |
| | Losartan | 0.11±0.16 | |
| tCompCor2 | Placebo | 0.03±0.14 | 0.07 |
| | Losartan | 0.02±0.17 | |
| tCompCor3 | Placebo | -0.03±0.20 | -0.74 |
| | Losartan | 0.00±0.11 | |
| tCompCor4 | Placebo | 0.06±0.22 | 1.81 |
| | Losartan | -0.06±0.30 | |
| tCompCor5 | Placebo | -0.07±0.46 | -1.71 |
| | Losartan | 0.16±0.56 | |

Abbreviation, tCompCor - temporal component based noise correction variant.

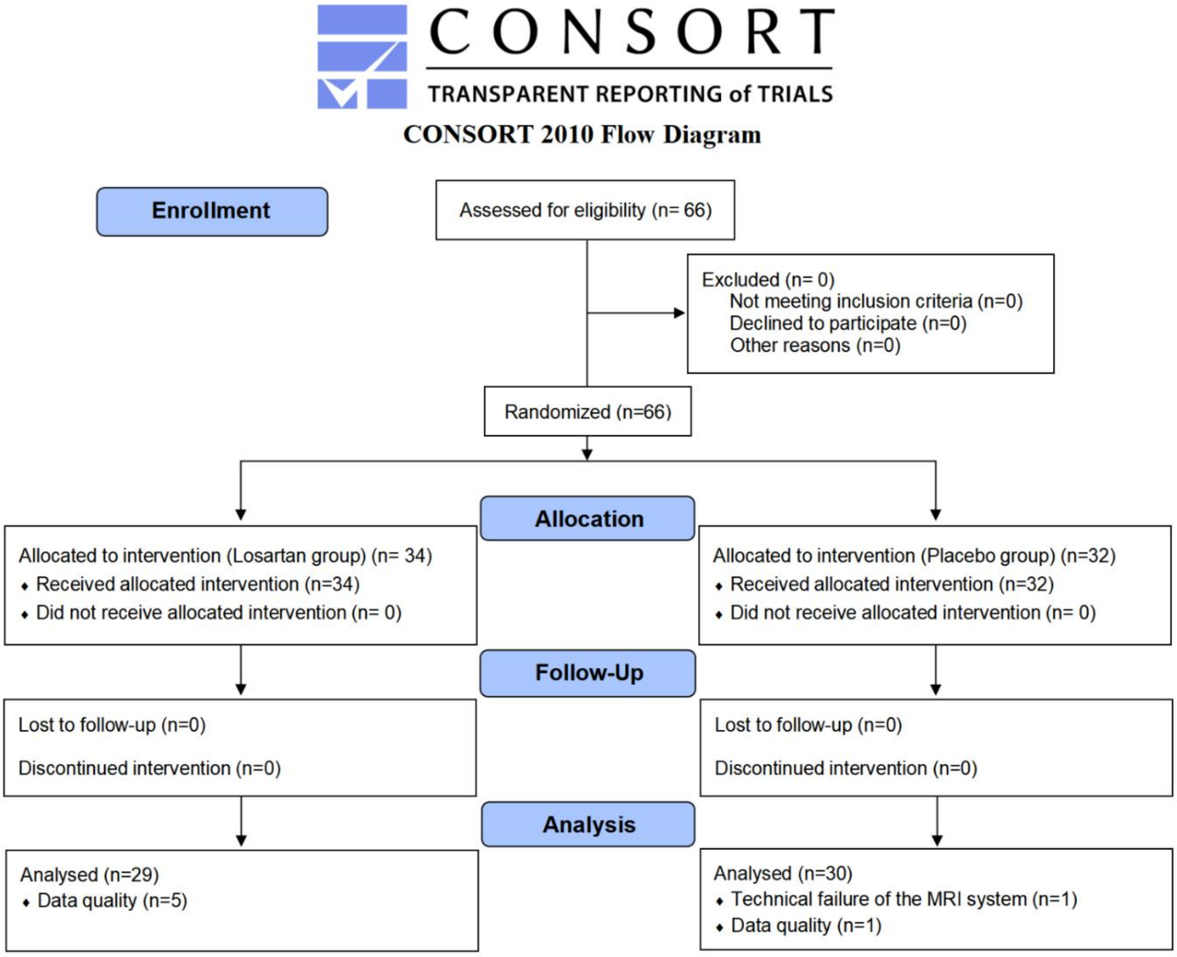


**Figure S1**. The CONSORT flow diagram

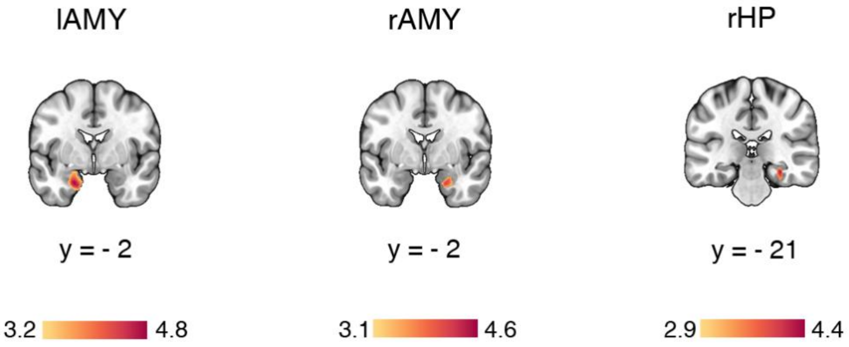


**Figure S2.** Regional activation associated with the successful memory formation. The color bar reflects the change of T value. lAMY, left amygdala; rAMY, right amygdala; rHP, right hippocampus.

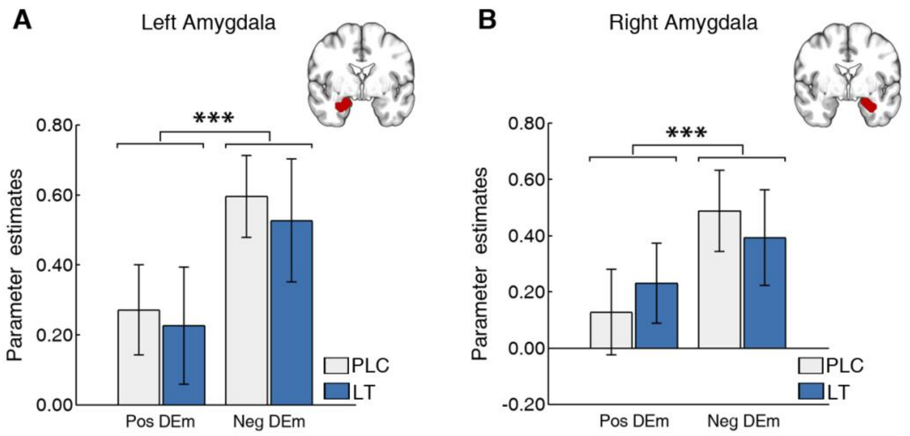


**Figure S3.** LT effects on neural activity during successful memory formation for emotional stimuli (DEm effect). **A-B** Bilateral amygdala displayed increased activation during memory formation for negative stimuli relative to positive ones irrespective of treatment. Error bars represent standard error of the mean. Pos, positive; Neg, negative; PLC, placebo; LT, losartan;***p<0.001.
